# Genome reduction occurred in early *Prochlorococcus* with an unusually low effective population size

**DOI:** 10.1101/2023.06.25.546417

**Authors:** Hao Zhang, Ferdi L. Hellweger, Haiwei Luo

## Abstract

In the oligotrophic sunlit ocean, the most abundant free-living planktonic bacterial lineages evolve convergently through genome reduction. The cyanobacterium *Prochlorococcus* responsible for 10% global oxygen production is a prominent example. The dominant theory known as ‘genome streamlining’ posits that they have extremely large effective population sizes (*N*_*e*_) such that selection for metabolic efficiency acts to drive genome reduction. Because genome reduction largely took place anciently, this theory builds on the assumption that their ancestors’ *N*_*e*_ was similarly large. Constraining *N*_*e*_ for ancient ancestors is challenging because experimental measurements of extinct organisms are impossible and alternatively reconstructing ancestral *N*_*e*_ with phylogenetic models gives large uncertainties. Here, we develop a new strategy that leverages agent-based modeling to simulate the change of *N*_*e*_ proxy for ancient ancestors, the genome-wide ratio of radical to conservative nonsynonymous nucleotide substitution rate (d_*R*_/d_*C*_), in response to the change of *N*_*e*_. Surprisingly, this proxy shows expected increases with decreases of *N*_*e*_ only when *N*_*e*_ falls to about 10k – 100k or lower, magnitudes characteristic of *N*_*e*_ of obligate endosymbiont species where drift drives genome reduction. We therefore conclude that drift, rather than selection, is the primary force that drove *Prochlorococcus* genome reduction.

## Main

Marine bacterioplankton cells in oligotrophic sunlit water are dominated by those that possess streamlined genomes ^1^. Small genomes confer a variety of fitness advantages to surface water bacterioplankton populations, including reduced biosynthetic requirement for nutrients which are exceedingly low in oligotrophic surface waters, increased rate of diffusive delivery of nutrients to the surface of the cells owing to concomitant cell size reduction and genome size reduction, and enhanced photon absorption in the case of photosynthetic bacteria ^2-4^. Because of these benefits, it has been widely accepted that the evolutionary transition from large genomes to small genomes is driven by positive selection, a process named ‘genome streamlining’ ^3^.

To make these advantages visible to selection, bacterioplankton lineages need to have sufficiently large effective population sizes (*N*_*e*_) so that natural selection overrides genetic drift (i.e., the inverse of *N*_*e*_), a random sampling effect that results in chance deletion of beneficial traits and chance fixation of deleterious variants. Since genome-reduced bacterioplankton lineages are often extremely abundant across the global ocean, they have been simply assumed to have very large *N*_*e*_ ^3^. However, a mismatch by several orders of magnitude between *N*_*e*_ and the census population size is common in nature due to demographic dynamics and selective sweeps among other evolutionary processes ^5,6^.

Estimating *N*_*e*_ of a bacterial species requires i) measuring its genomic mutation rate with an approximately unbiased method, commonly through a mutation accumulation (MA) experiment followed by whole-genome sequencing (WGS) of derived lines, and ii) calculating the genetic diversity at neutral sites (usually approximated by synonymous sites) across multiple strains capturing the diversity of the species ^7^. Such data are rarely available for genome-reduced marine bacterioplankton lineages, since the requirement of propagating the bacterial isolates of these lineages on solid media in many parallel lines across hundreds of generations in a MA experiment is difficult to meet. To date, the *Prochlorococcus* high light-adapted clade II (HLII) has been the only genome-reduced bacterioplankton lineage whose *N*_*e*_ was estimated through this approach, and the same type of data is available for more large-genome marine bacterial lineages (mainly *Ruegeria* and *Vibrio*) ^8^. Apparently, the genome streamlining theory implies that genome-reduced bacterioplankton lineages have greater *N*_*e*_ than those with large genomes. Therefore, the surprising finding ^8^ that *N*_*e*_ of HLII (1.68 × 10^7^) is one order of magnitude smaller than that of a *Ruegeria* species (1.85 × 10^8^) and three *Vibrio* species (1.12∼2.21 × 10^8^) is not consistent with the genome streamlining theory, though this conclusion is based on limited taxonomic coverage.

Another major problem is that arguments for genome streamlining often inadvertently build on the presumably large *N*_*e*_ of extant species, ignoring the fact that genome reduction often occurred in the distant past. In the case of *Prochlorococcus* and SAR11, the two most abundant marine bacterioplankton lineages, genome reduction largely occurred in their early evolution ^9-11^, far beyond the evolutionary timescale spanned by any extant species. Hence, to gain further insights, it is essential to constrain the *N*_*e*_ value at the ancient time when genome reduction occurred, which is challenging for all species, particularly the non-fossilized species such as marine bacterioplankton lineages.

*Prochlorococcus* are the most abundant photosynthetic carbon-fixing organisms in the modern ocean, responsible for ∼10% of marine net primary production ^12^. *Prochlorococcus* have evolved multiple ecotypes representing ecologically distinct and evolutionarily separate lineages, which are broadly distinguished between HL adapted ecotypes and low light (LL)- adapted ecotypes ^13^. The HL ecotypes (HLI and HLII among others) form a monophyletic group, whereas the LL ecotypes (LLI, LLII/III, and LLIV among others) form a paraphyletic group where the HL ecotypes are rooted (Fig. 1A) ^13^. Following the divergence of the basal LLIV, *Prochlorococcus* experienced major genome reduction, resulting in reduced genomes carried by LLII/III, LLI, and HL ecotypes compared to those by LLIV ^2,10,11,14^.

**Fig. 1.**
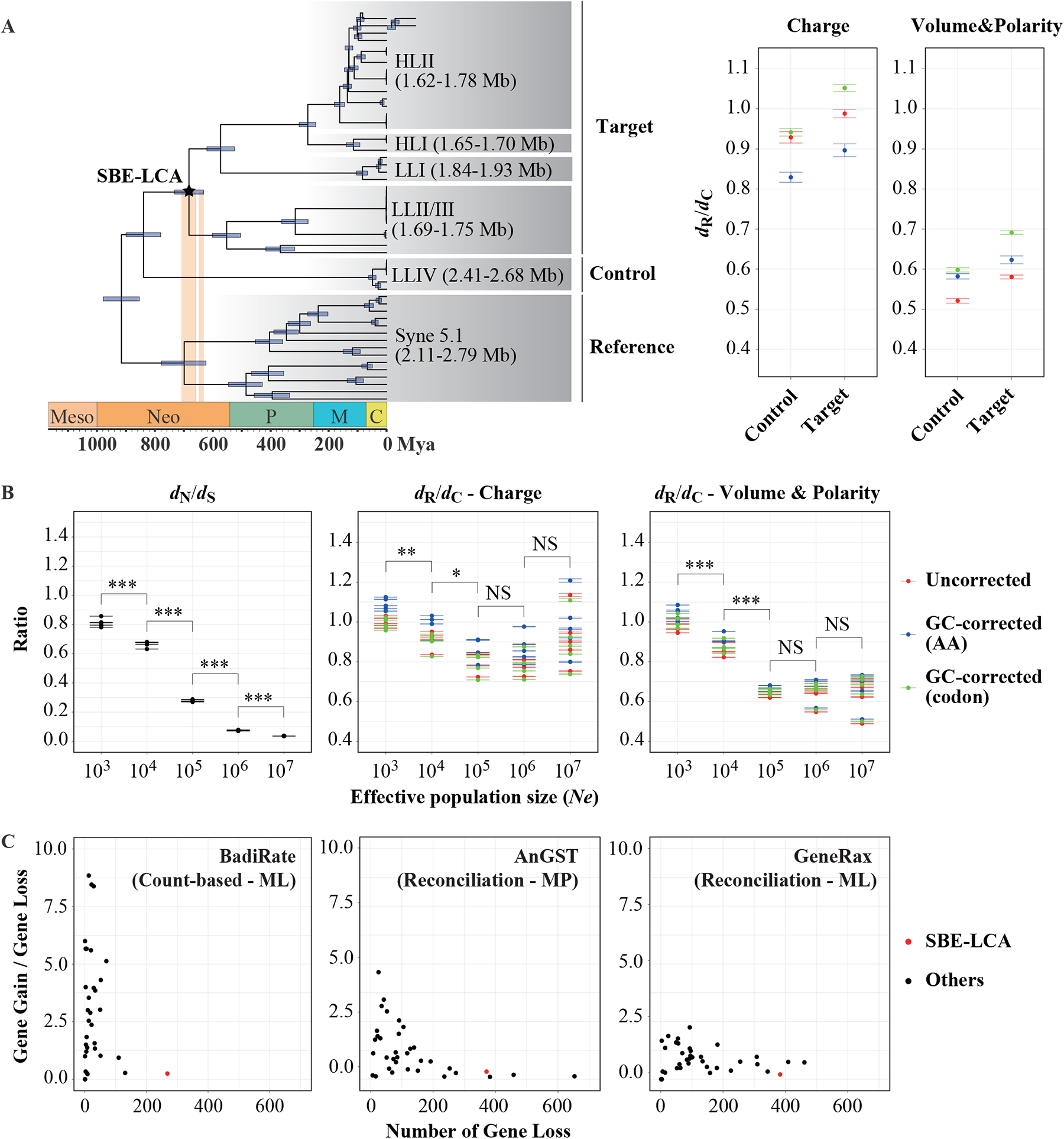
The *Prochlorococcus* evolutionary timescale, mechanism and process during their early evolution when a major genome reduction event occurred. (A) The chronogram of *Prochlorococcus* evolutionary timeline adapted from Zhang et al. (2021). The left and right blue vertical bars represent the Sturtian and Marinoan glaciations, respectively, during the Neoproterozoic Snowball Earth. The blue horizontal bar on each ancestral node represents the posterior 95% highest probability density (HPD) interval of the divergence time estimated by MCMCTree v4.10.0. The ancestral node (SBE-LCA) marked with star experienced severe population bottleneck. The right panel shows a greater genome-wide *d*_R_/*d*_C_ ratio of the monophyletic group derived from the pairwise comparisons between genomes from the target group (all descendants of SBE-LCA) and genomes from the reference group (the *Synechococcus* cluster 5.1) than that derived from the pairwise comparisons between genomes from the control group (*Prochlorococcus* LLIV clade) and genomes from the reference group. Note that these figures are adapted from Zhang et al., 2021. (B) The genome-wide *d*_N_/*d*_S_ and *d*_R_/*d*_C_ ratios of simulated *Prochlorococcus* populations under different effective population sizes (*Ne*). For each *Ne*, the simulation ran 10 times and each generated 50 genome sequences. Mean values of the *d*_N_/*d*_S_ and *d*_R_/*d*_C_ ratios in each run are used for one-tailed t-test (***: p<0.001, **: p<0.01, *: p<0.05; NS: non-significant). The GC-corrected *d*_R_/*d*_C_ values based on codon frequency and AA composition are marked in red and blue, respectively, and the uncorrected *d*_R_/*d*_C_ values are marked in green. The error bars in the plots represent the standard error of the mean (SEM). (C) The number of gene loss events and the ratio of gene gain to loss events reconstructed with BadiRate (adapted from Zhang et al., 2021), AnGST, and GeneRax. In each plot, the dots represent the ancestral nodes shown in (A), among which the SBE-LCA is marked in red. Ancestral nodes with extremely high gene gain to gene loss ratios (5, 4, and 1, respectively, in the panel labeled with BadiRate, AnGST and GeneRax) were detected with the boxplot.stats function in R and eliminated from the plots.

Recently, we reconstructed a timeline of *Prochlorococcus* evolution and showed that the main genome reduction took place during the Neoproterozoic Snowball Earth (Fig. 1A; denoted as “SBE-LCA”) ^15^. This conclusion remains robust after accounting for the uncertainty of fossil calibrations and root age constraints and optimizing a variety of parameters with statistical tests such as choosing alternative molecular clock models. Since *Prochlorococcus* are generally adapted to warm waters, as shown by their ecological distribution restricted to low latitudes between 40 °N and 40 °S ^13^ and by their physiology that all tested HL and LL isolates rarely grow at temperatures below ∼10°C ^16^, the global icehouse climate conditions during the Snowball Earth ^17,18^ likely drove population bottlenecks, which left strong signatures in genomic DNA through the accumulation of deleterious mutations ^15,19^.

To put deleterious mutations in context, we distinguished two types of nonsynonymous mutations – radical and conservative changes leading to chemically dissimilar and similar amino acid replacements, respectively, and we used *d*_R_ and *d*_C_ to represent the rates of radical and conservative changes, respectively. We first provided evidence that radical changes are under stronger purifying selection and thus more deleterious than conservative changes in both HL and LL *Prochlorococcus* lineages ^19^. This is done by dividing nonsynonymous SNPs of intraspecific *Prochlorococcus* genomes into chemically similar and dissimilar SNP groups according to their encoded amino acids, followed by comparing heterozygosity between the two groups. According to theory, the SNP group that displays lower heterozygosity is under more intensive purifying selection, thereby being more deleterious. We next showed an excess of radical changes relative to conservative changes across the genomic regions (i.e., an inflated genome-wide *d*_R_/*d*_C_ ratio) in the monophyletic group comprising LLII/III, LLI, and HL ecotypes compared to that in LLIV. This is evidence that genetic drift acted at the last common ancestor (i.e., SBE-LCA) of LLII/III, LLI, and HL ecotypes, coinciding with the major genome reduction of *Prochlorococcus* (Fig. 1A) ^15,19^.

This coincidence does not necessarily mean that genetic drift acted to drive genome reduction. This is because the *N*_*e*_ value at that ancestral stage is not known. If the population bottlenecks were weak and *N*_*e*_ remained large, selection may have remained efficient and thus could still be the primary driver of genome reduction. Therefore, constraining the *N*_*e*_ value of the *Prochlorococcus* populations during the Neoproterozoic Snowball Earth becomes the key step to disentangle the evolutionary mechanisms underlying *Prochlorococcus* genome reduction. For such ancient and thus extinct populations, one cannot extrapolate their *N*_*e*_ values by performing a MA experiment and collecting multiple strains for calculating genetic diversity at neutral sites. Instead, we develop an agent-based model (ABM) to constrain the *N*_*e*_ values by linking *N*_*e*_ to the *d*_R_/*d*_C_ ratio.

Our model builds on a previous ABM we developed to study the genomic G+C content evolution of a heterotrophic marine bacterium *Ruegeria pomeroyi* DSS-3 ^20^. The model simulates a population of individual cells, each with a complete genome, that grow, divide, mutate and recombine. Here we consider functional changes of mutations, i.e. effects on carbon and nitrogen requirements of DNA and protein pools are ignored. The model was modified to explicitly consider deleterious and advantageous nonsynonymous mutations. For each mutation, a selection coefficient is drawn randomly from a probability distribution with negative (deleterious) and positive (advantageous) values, where the magnitude is a function of the chemical distance of the corresponding amino acids. This coefficient is then used to scale the growth rate of the individual. The mutation is inherited by offspring and over subsequent generations affects the success of the mutation in the population, and with that the model can simulate the effect of purifying selection (Fig. S1; see details in SI)

To validate the new model, we tracked the change of the *d*_N_/*d*_S_ ratio following the change of the *N*_*e*_ values. The *d*_N_/*d*_S_ ratio is the rate of nonsynonymous nucleotide substitution relative to the rate of synonymous nucleotide substitution. Because synonymous substitutions are largely neutral and nonsynonymous substitutions are more likely to be deleterious, an elevated genome-wide *d*_N_/*d*_S_ is broadly used as a proxy for an increased power of genetic drift because of reduced *N*_*e*_ ^21^. Mathematically, the *d*_N_/*d*_S_ ratio gives a composite parameter, *N*_*e*_*s* ^22^, where *s* is the selection coefficient of the genes under the analysis and may vary over time and between lineages. In our simulation, the *d*_N_/*d*_S_ ratio was calculated based on the same genes from the same lineage and it was compared between simulations under different *N*_*e*_ values. Thus, *s* is a constant and *d*_N_/*d*_S_ must show a monotonical decrease with increasing *N*_*e*_ if our model works well, which we have demonstrated in Fig. 1B. We then calculated the *d*_R_/*d*_C_ ratio using the same simulated data and surprisingly found that *d*_R_/*d*_C_ decreases with increasing *N*_*e*_ only when *N*_*e*_ is on the order of 10^4^-10^5^ or below, and becomes statistically invariable with further increases of *N*_*e*_ (Fig. 1B). This finding is robust to the changes of growth rate, a parameter likely differentiating between ancient and modern *Prochlorococcus*, employed in simulations (Fig. S2). We further benchmarked our model with three other phylogenetically diverse bacterial species, and consistently found that *N*_*e*_ on the order of 10^4^- 10^5^ as the threshold that differentiates the response of *d*_R_/*d*_C_ to the change of *N*_*e*_ (Fig. S3).

Our simulation results indicate that whereas population bottlenecks (e.g., a decrease of *N*_*e*_ from 10^7^ to 10^5^) do not necessarily lead to an increase of *d*_R_/*d*_C_, an increase of *d*_R_/*d*_C_ can be linked to population bottlenecks that decrease *N*_*e*_ to the order of 10^4^-10^5^ or lower. Hence, while the genome-wide *d*_R_/*d*_C_ ratio is not a sensitive proxy for the change of *N*_*e*_, it constrains the magnitude of *N*_*e*_ if it changes. Notably, *N*_*e*_ on the order of 10^5^ matches that of obligate endosymbiotic bacterial species. For example, *Mesoplasma florum*, the only obligate endosymbiotic species with its *N*_*e*_ calculated based on its global mutation rate determined through the MA/WGS strategy ^23^ and genetic diversity at synonymous sites has *N*_*e*_ of 7.88 × 10^5 8^. The *N*_*e*_ estimates of other obligate endosymbionts are also available, such as *Buchnera* species which are on the order of 10^7 24^, but the mutation rate used for *N*_*e*_ estimation was derived from comparative sequence analyses ^25^ and thus less accurate. Because obligate endosymbionts are paradigms for genetic drift as the primary driver of bacterial genome reduction ^11,26,27^, the comparable *N*_*e*_ values between bottlenecked *Prochlorococcus* during the Neoproterozoic Snowball Earth and obligate endosymbiotic species indicate that genetic drift also acted to drive *Prochlorococcus* genome reduction.

To play a part in genome reduction, genetic drift needs to couple with mutational processes. One important consequence of genetic drift is increased mutation rate. According to the ‘drift-barrier’ model, natural selection favors low mutation rate and thus promotes replication fidelity to a threshold set by genetic drift, and further refinements are expected to decrease fitness advantages ^23^. This model explains well the mutation rate variation among prokaryotic species ^8^ and across prokaryotes and eukaryotes ^7^. Theory predicts that when mutation rate exceeds a certain threshold, selection cannot maintain all non-essential genes ^28^. For instance, a factor of 10 fold increase in mutation rate is predicted to lead to a 30% decrease in genome size by a mathematical model ^28^. In fact, a correlation between an increased mutation rate and genome reduction was recently established for *Prochlorococcus* through a phylogenetic analysis ^29^. It is worth mentioning that HLII, one of the most genome-reduced *Prochlorococcus* lineages, has a low genomic mutation rate (3.50 × 10^−10^ per site per generation) determined by the MA/WGS strategy, close to that of six other surface ocean bacterial species, including one *Ruegeria* species in Alphaproteobacteria (1.39 × 10^−10^) ^30^, three *Vibrio* species in Gammaproteobacteria (1.07, 2.07, 2.29 × 10^−10^) ^31,32^, and one *Nonlabens* species and one *Leeuwenhoekiella* species in Flavobacteriia (5.21, 5.49 × 10^−10^) ^33^. Therefore, it remains possible that drift and increased mutation rate acted together to reduce *Prochlorococcus* genome size during the Neoproterozoic Snowball Earth, but this hypothesis does not gain full support from experimental measurement of the modern *Prochlorococcus* mutation rate.

Alternatively, restricted opportunities for gene exchange may exacerbate the effect of genetic drift on genome reduction. In a small population with limited recombination, deleterious mutations are accumulated by random loss of the genotypes carrying least deleterious mutations and by random increase of the genotypes harboring more deleterious mutations, and they can only be removed by back mutations which occur very infrequently ^26^. As a result, non-essential genes gradually become pseudogenes and are subsequently deleted because of the deletional bias of the mutational spectrum commonly found in bacteria ^34^ including *Prochlorococcus* as determined by the MA/WGS strategy ^8^, and the deleted genes can hardly be regained because of limited recombination ^11^. This process is known as ‘Muller’s ratchet’, which is the major mechanism of genome degradation in obligate endosymbionts ^26,35^, but is usually limited in explaining genome reduction in free-living bacteria which are generally believed to have large population sizes and engage freely in recombination ^36^. During the Neoproterozoic Snowball Earth, however, *Prochlorococcus* had very small *N*_*e*_ at the order of 10^4^-10^5^ or lower as shown here, which is comparable to obligate endosymbionts. Further, cells were not connected but instead were restricted to a few biotic refugia mainly including sea-ice brine channels and cryoconite holes ^15,37,38^, which may have appreciably reduced the recombination rate.

To investigate the recombination frequency of *Prochlorococcus* in the ancestral branches, we reconstructed gene gains and losses along the species phylogeny and used gene gain/loss ratio of each ancestral branch as a proxy. We adapted the previously reconstructed genomic events from our recent study ^15^, benchmarked a few other methods in common use (see details in SI; Fig. S4), and consistently found that the genome content changes of *Prochlorococcus* during the Neoproterozoic Snowball Earth were dominated by gene loss and that the gene gain/loss ratio approached to zero and is among the smallest across all phylogenetic branches in *Prochlorococcus* phylogeny (Fig. 1C). Since these results provide evidence for low recombination rate of *Prochlorococcus* during the time when major genome reduction occurred, we propose that population bottlenecks and rare recombination together may have escalated the effect of Muller’s ratchet on *Prochlorococcus* genome reduction.

The available *Prochlorococcus* isolates represent snapshots of the genome eroding process, but how genetic drift causes genome reduction is not clear. In genome-reduced symbionts like *Buchnera*, it was proposed that genetic drift led to the losses of DNA repair genes and a subsequent shift of genomic nucleotide composition toward high A+T content. Genomes with A+T enrichment have an excess of A/T homopolymers and a resulting high incidence of small indels leading to increased gene inactivation ^39^. Since all genome-reduced *Prochlorococcus* ecotypes (LLII/III, LLI, and HL) have higher A+T content than the basal LLIV ecotype (62-69% versus 49-50%) and the former have fewer DNA repair genes than the latter ^2^, it is reasonable to hypothesize that the above model theorized for symbionts may also fit *Prochlorococcus*. Alternatively, an increased power of genetic drift may allow expansion of mobile genetic elements (MGEs) whose transpositions lead to gene inactivation ^40^. This mechanism was proposed for symbiont genome reduction but remains unknown for *Prochlorococcus* and other genome-reduced marine bacterioplankton lineages in which MGEs are overall rare.

Although *Prochlorococcus* represents a paradigm for the study of the processes and mechanisms underlying marine bacterioplankton genome reduction, it is not clear whether the mechanisms learnt from *Prochlorococcus* are equally important to drive genome reduction of other marine bacterioplankton lineages. Marine SAR11 is another classical lineage that argues for streamlining selection ^41^. Marine SAR11 genomes have strong signatures of recombination and have some of the highest r/m ratios (measuring the relative effect of recombination to mutation on genetic diversity of a bacterial species) among all known marine and non-marine bacterial lineages, whereas the *Prochlorococcus* populations have very small ratios (61-63 versus 1-3) ^8,42-44^. Since recombination is the main path that eliminates deleterious mutations, high recombination rate increases selection efficiency. Therefore, the extant marine SAR11 lineages likely have very different genetic population structure than the extant *Prochlorococcus* lineages, and a possibility of a greater role of selection on SAR11 evolution than on *Prochlorococcus* evolution cannot be ruled out.

While the key drivers triggering genome reduction in most marine bacterioplankton lineages remain debated, consequences of genome reduction have already become clear. Cells may benefit from harboring small genomes by reduced requirements of nutrients for biosynthesis and enhanced efficiency of nutrient (and light) assimilation. These fitness advantages may be sufficiently large for modern bacterioplankton populations such that small genomes are maintained by purifying selection. In a warming and more stratified future ocean, nutrients will become increasingly limited to surface ocean bacterioplankton cells. Under this stronger selective constraint, it can be predicted that smaller genomes will confer greater fitness advantage to surface ocean bacterioplankton populations. As such, pelagic microbial populations that have already carried small genomes, regardless of the evolutionary paths by which the genomes evolved to become small, will either stay as what they are or become further reduced by deleting unnecessary DNA, and expansions of their genomes will be selectively disfavored in an increasingly warming and stratified ocean.

The exact evolutionary mechanisms that acted to drive genome reduction and evolution in the ocean are relevant to the biogeochemical processes that marine bacterioplankton lineages mediate. During the major *Prochlorococcus* genome reduction stage, some beneficial traits likely including ecologically relevant genes were expectedly lost because of the escalated power of genetic drift. Moreover, the remaining genetic traits in *Prochlorococcus* should no longer be considered fully optimized by natural selection, but instead represent a genetic repertoire embedded with an excess of mildly deleterious mutants. If the genes that encode key environmental functions are enriched with deleterious mutants, the related biochemical processes such as photosynthesis, carbon fixation, and nutrient acquisition may not be as efficient as expected. Given the global dominance of *Prochlorococcus*, the compromised efficiency is relevant to the carbon, nitrogen and other elemental cycling in the ocean. An improved understanding of the evolutionary drivers of genome-reduced bacterioplankton lineages has important implications for the role of pelagic microbial community in climate regulation.

## Data and code availability

The source code and sample files for ABM simulation, as well as the custom scripts for genome-wide *d*_N_/*d*_S_ and *d*_R_/*d*_C_ calculation are deposited in the online GitHub repository (https://github.com/luolab-cuhk/Prochl-ABM).

## Acknowledgements

This work is supported by the National Natural Science Foundation of China (42293294), the Hong Kong Research Grants Council Area of Excellence Scheme (AoE/M-403/16), the Natural Science Foundation of Guangdong Province, China (2022A1515010844 to HZ), and the China Postdoctoral Science Foundation (2021M702296 to HZ).

