## Supplementary material for "Genome reduction occurred in early *Prochlorococcus* with an unusually low effective population size": SI

**This file includes:**

**Methods**

**References**

**Fig. S1 to S4**

### Methods

#### Agent-based model building

We used an updated version of the IAM model presented previously (Hellweger et al., 2018). The model code and sample input files are deposited in the online GitHub repository (https://github.com/luolab-cuhk/Prochl-ABM).

To support the simulations presented here, an additional nonsynonymous mutation penalty sub-model (no. 8, see Table S1 in Hellweger et al., 2018 and associated text and references) was implemented in the code. This was done for the following reason. In the previous version, nonsynonymous mutations were assumed to be either functionally neutral or deleterious, with the probability of being deleterious dependent on the amino acid chemical distance. Deleterious mutations were assumed to be lethal, which was done to exclude them from the population, otherwise they would gradually lower the population growth rate (i.e. Muller's Ratchet). This approach is realistic for constant and large effective population size ($N_e$), as was the case in the previous model application. Note that the previous application focused on the role of nutrient limitation in shaping the genome, and not the effect of $N_e$ on $d_N/d_S$ or $d_R/d_C$. However, this approach does not explicitly simulate purifying selection and the effect of $N_e$ on that, which is the focus of the present model application. Here, the deleterious mutations have to "stay in the population" (i.e., we cannot simply remove them by making the mutation fatal) so that they can be outcompeted, which will be a function of $N_e$.

The new nonsynonymous mutation fitness (previously called penalty) sub-model is as follows. For each nonsynonymous mutation, a selection coefficient ($s$) is drawn from a probability distribution. The distribution, illustrated in Fig. S1, consists of an exponential distribution below 0 (deleterious mutations) and a uniform distribution above 0 (advantageous mutations). The distribution is defined by three parameters, including the fraction of mutations that are deleterious ($f_{del}$), and the average selection coefficients for the deleterious mutations ($s_{pn}$) and the advantageous mutations ($s_{an}$). $f_{del}$ and $s_{an}$ are global parameters. $s_{pn}$ is a function of the amino acid (AA) dissimilarity. Specifically, the $s_{pn}$ for mutations between AA $i$ and $j$ is:

$$s_{pn}(i,j) = s_{base}\frac{d(i,j)}{d_{base}}$$

$s_{pn} = s_{base}$ for $d(i,j) = d_{base}$.

$d(i,j)$ = AA dissimilarity matrix.

The selection coefficients for all occurred mutations affect the growth rate (via the nutrient uptake rate, see previous model description). The model does not explicitly consider any biological constraints on the overall fitness of a cell and to avoid biologically unrealistic increases in fitness (i.e., Darwinian Demon), the growth rate is not allowed to exceed (i.e., is capped at) that of the ancestor in the simulation.

For recombination, selection coefficients of incoming changes/mutations from the donor overwrite those of the recipient in the simulation.

**Agent-based model application**

Four bacterial species were chosen as the targets for modelling with ABM, including *Prochlorococcus marinus* AS9601 (GCF_000015645), *Bacillus subtilis* NCIB 3610 (GCF_002055965), *Ruegeria pomeroyi* DSS-3 (GCF_000011965), and *Vibrio fischeri* ES114

(GCF_000011805). They each have publicly available data of unbiased global mutation rate determined by the MA/WGS strategy ($3.50 \times 10^{-10}$, $3.28 \times 10^{-10}$, $1.39 \times 10^{-10}$, and $2.07 \times 10^{-10}$ per site per generation, respectively) (Sung et al., 2016; Dillon et al., 2017; Sun et al., 2017; Chen et al., 2022) and they are phylogenetically diverse. Their genome sequences were downloaded from the curated NCBI RefSeq database. We employed Prodigal v2.6.3 (Hyatt et al., 2010) to call protein-coding genes on genomic sequence and then generate the genomic feature file (GFF). GFF records the start and end positions of ORFs and was used as the input for genome simulation (see example file in the online GitHub repository).

In our simulations, $N_e$ is a function of the census population size ($N_c$) at the beginning of the growth period ($N_c0$) and the number of generations in the dilution step ($n_G$) as follows:

$$N_e = N_c0 * n_G$$

The factor $n_G$ is another function of the growth rate [$k_G$, determined by the availability of C ($V_{max}0C$), N ($V_{max}0N$), and P ($V_{max}0P$)] and the dilution time step ($d_t$) as follows:

$$n_G = \frac{d_t}{(\frac{ln2}{k_G})}$$

All the abovementioned model parameters are deposited in a configuration file, which is required for our simulation (see the example configuration file in online GitHub repository). Note that the default parameters for the nonsynonymous mutation fitness model were calibrated for *Prochlorococcus marinus* AS9601, so that $d_N/d_S = 0.05$ for $N_e = 10^7$. We ran each simulation five times with different seed number ($RS_i$ and $RS_j$ in the configuration file) to reduce the random error and perform statistical tests. As each run generates 50 genome sequences, we obtained a total of $50 \times 5$ genome sequences under each putative $N_e$. To simulate the genome sequences under different growth rates, we revised the growth rate-associated parameters ($V_{max}0C$, $V_{max}0N$ and $V_{max}0P$) to 10%, 50% and 200% of the default. According to above equations, we revised the dilution time step ($d_t$) to 1000%, 200% and 50% of the default, respectively, to keep a constant $N_e$ in these simulations. Note that for 10% growth rate simulations, we also increased the simulation time (*tend* in the configuration file) to ensure accumulating a sufficient number of substitutions in their genome sequences. Whereas the model includes various effects of mutations, e.g. changes to the C, N, and P requirements of DNA and protein pool, the present application only considers the effect of nonsynonymous mutations on protein function.

**Genome-wide $d_N/d_S$ and $d_R/d_C$ calculation**

We extracted protein-coding gene sequences from simulated genomes with in-house scripts based on GFF instead of using Prodigal to avoid skipping ORFs in which substitutions occurred at start or stop codon. We estimated the $d_N$ and $d_S$ values of each protein-coding gene with the program YN00 in PAML package v4.9e (Yang and Nielsen, 2000), which takes into account of the transition/transversion ratio (ts/tv) bias and the codon usage bias. To obtain the genome-wide $d_N/d_S$ value, let $N_{i(x)}$ and $S_{i(x)}$ be the number of nonsynonymous and synonymous substitutions and let $N_{j(x)}$ and $S_{j(x)}$ be the nonsynonymous and synonymous sites for each gene ($x$). Let $n$ be the number of genes in the focal bacterial strain. The genome-wide $d_N/d_S$ is computed as:

$$d_N/d_S = \left(\frac{\sum_1^n N_{i(x)}}{\sum_1^n N_{j(x)}}\right)/\left(\frac{\sum_1^n S_{i(x)}}{\sum_1^n S_{j(x)}}\right)$$

To estimate the $d_R$ and $d_C$ values for each gene, we employed the software MEGA-CC v10.2.4 (Kumar et al., 2008) to calculate ts/tv and then passed the ratio to the software RCCalculator (Luo et al., 2017). The latter recruits two new models, either based on amino acid frequency or based on codon frequency, to correct for G+C content bias. To obtain the genome-wide $d_R/d_C$ value, let $R_{i(x)}$ and $C_{i(x)}$ be the number of radical and conservative nonsynonymous substitutions and let $R_{j(x)}$ and $C_{j(x)}$ be the radical and conservative nonsynonymous sites for each gene ($x$). Let $n$ be the number of genes in the focal bacterial strain. The genome-wide $d_R/d_C$ is computed as:

$$d_R/d_C = (\frac{\sum_1^n R_{i(x)}}{\sum_1^n R_{j(x)}})/(\frac{\sum_1^n C_{i(x)}}{\sum_1^n C_{j(x)}})$$

**Inferring gene gain and loss events along species phylogeny**

The gene gain and loss process along phylogeny is commonly reconstructed by reconciliating incongruence between gene tree and species tree. There are multiple reconciliation software tools, which often give different predictions. To find out the tool that fits best our dataset, we benchmarked four tools in common use, including the likelihood-based ALE v0.4 (Szöllősi et al., 2013) and GeneRax v2.0.4 (Morel et al., 2020), as well as the parsimony-based AnGST v1.0 (David and Alm, 2011) and ecceTERA v1.2.4 (Jacox et al., 2016).

Our benchmarking strategy requires user-specified gene tree as “real tree”, uses a tool to simulate sequence alignment from the “real tree”, constructs initial gene tree from the simulated alignment and generates a reconciled gene tree, and employs the Robinson-Foulds

(RF) distance to assess the topological difference between the reconciled tree and the "real tree" (Fig. S4). To reduce the computational cost while keeping the complexity of the input gene tree, we compiled a small-scale dataset containing 1,000 out of the 4,689 pre-identified gene families of *Prochlorococcus* (Zhang et al., 2021) by sorting the gene family size and sampling with a fixed periodic interval (family size interval=4). Since true gene tree for empirical data is not available and since gene tree reconstructed by using reconciliation method is often more accurate than that reconstructed based on molecular sequence alone (Szöllősi et al., 2013), we reconciled the IQ-TREE-derived gene tree with *Prochlorococcus* species tree using AnGST v1.0 and GeneRax v2.0.4 to generate the "real tree" dataset I and the "real tree" dataset II, respectively (Fig. S4). The implementation of the two datasets helps to reduce the bias in benchmarking analysis towards either the maximum likelihood (ML) or the maximum parsimony (MP) algorithm. Note that the *Prochlorococcus* species tree and gene family tree used in this step were all adapted from our recent study (Zhang et al., 2021). For each "real tree", we employed the tool Bppseqgen v2.4.0 (Dutheil and Boussau, 2008) to simulate a sequence alignment under LG amino acid substitution model with Gamma-distributed 10% across-site rate variation. Simulated alignments were then subjected to IQ-TREE v2.0.6 for phylogeny inference with LG+G model and 1,000 ultrafast bootstraps. The bootstrapped trees were then reconciled with the *Prochlorococcus* species tree using the tool ALE v0.4, AnGST v1.0 and ecceTERA v1.2.4, while the IQ-TREE-derived ML tree was used as the initial tree for GeneRax reconciliation. Note that for parsimony-based reconciliations, the costs for gene duplication, gene transfer and gene loss (DTL) events were set to 3, 2 and 1, respectively, which are default values for many parsimony-based reconciliation tools (David and Alm, 2011; Bansal et al., 2015; Morel et al., 2020) and have been applied in several studies (Nagy et al., 2014; Hehemann et al., 2016). The qualities of these reconciled gene trees were measured by using the “RF.dist” function in the “phangorn” R package. We found that, for both “real tree” datasets, AnGST had lower mean RF distances and thus outperformed ecceTERA (for MP tools) and GeneRax outperformed ALE (for ML tools) (Fig. S4). We therefore applied AnGST and GeneRax to reconcile the full set of 4,689 *Prochlorococcus* gene family tree with the species tree.

**Fig. S1**

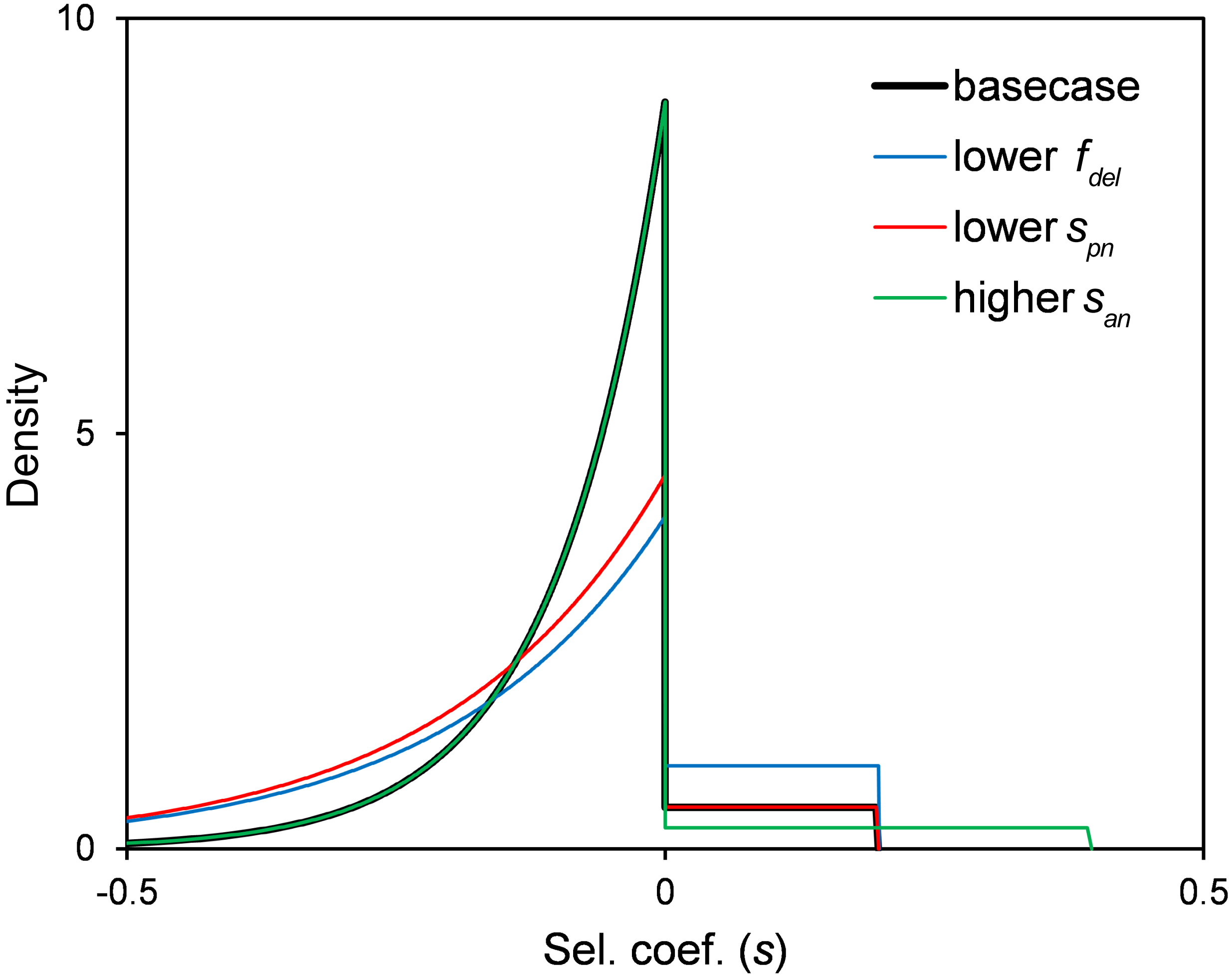


Fig. S1 (A) Nonsynonymous mutation fitness model. Parameters for basecase: $f_{del} = 0.9$, $s_{pn} = -0.1$, $s_{an} = 0.1$. Mutations with larger amino acid chemical distance will have lower $s_{pn}$ values (see details in SI and in Hellweger et al., 2018).

**Fig. S2**

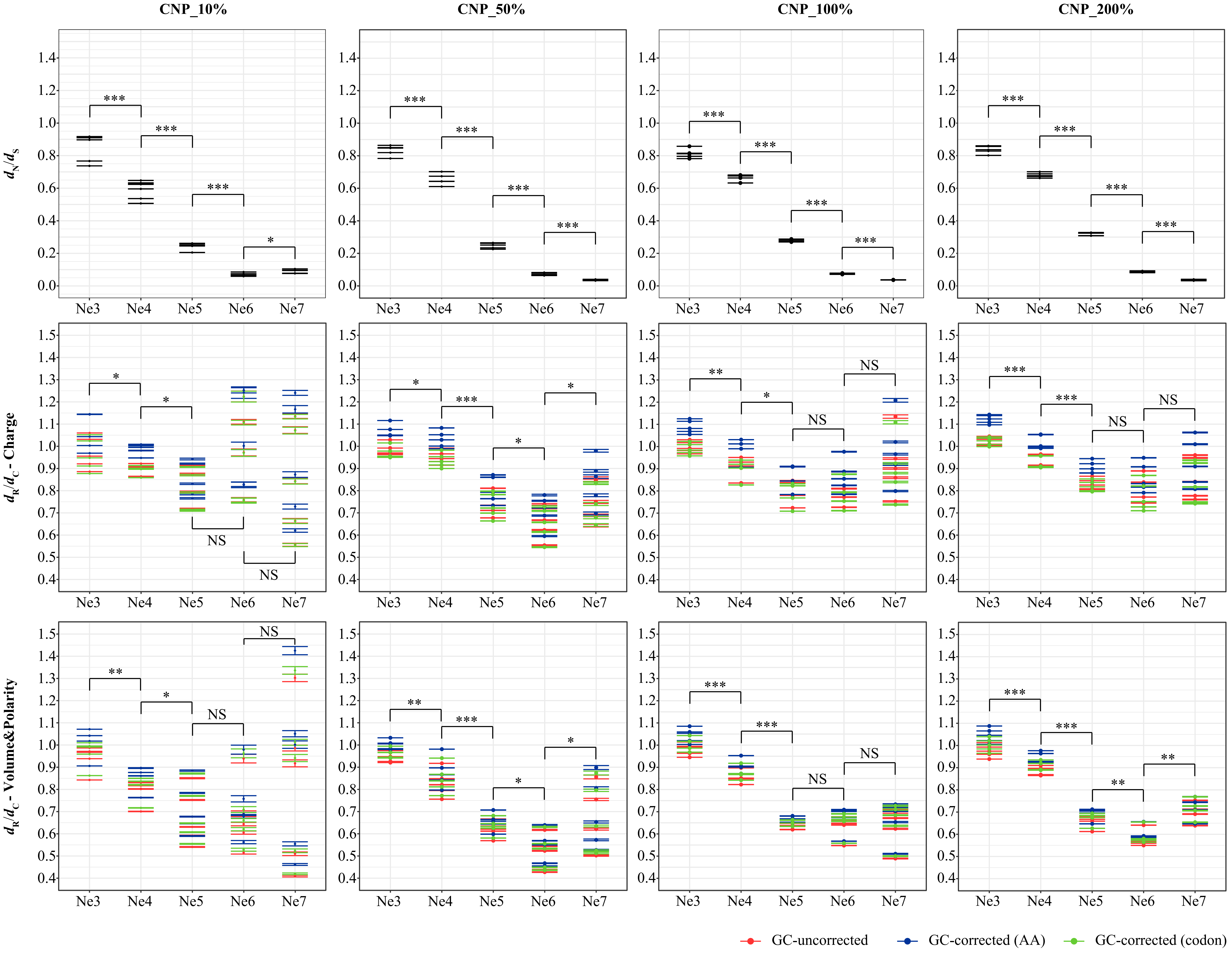


Fig. S2 The genome-wide $d_N/d_S$ and $d_R/d_C$ ratios of simulated *Prochlorococcus* populations under different growth rates. As bacterial growth rate is associated with the concentration of nutrient C, N and P, we controlled the nutrient-associated parameters in our simulations and used CNP_10%, CNP_50% and CNP_200% to represent the tenth, the half and the double growth rate of *Prochlorococcus*. For each $N_e$, the simulation ran five times and each generated 50 genome sequences. Mean values of the $d_N/d_S$ and $d_R/d_C$ ratios in each run are used for one-tailed t-test (***: $p<0.001$, **: $p<0.01$, *: $p<0.05$; NS: non-significant). The GC-corrected $d_R/d_C$ values based on codon frequency and AA composition are marked in red and blue, and the uncorrected $d_R/d_C$ values are marked in green. The error bars in the plots represent the standard error of the mean (SEM).

**Fig. S3**

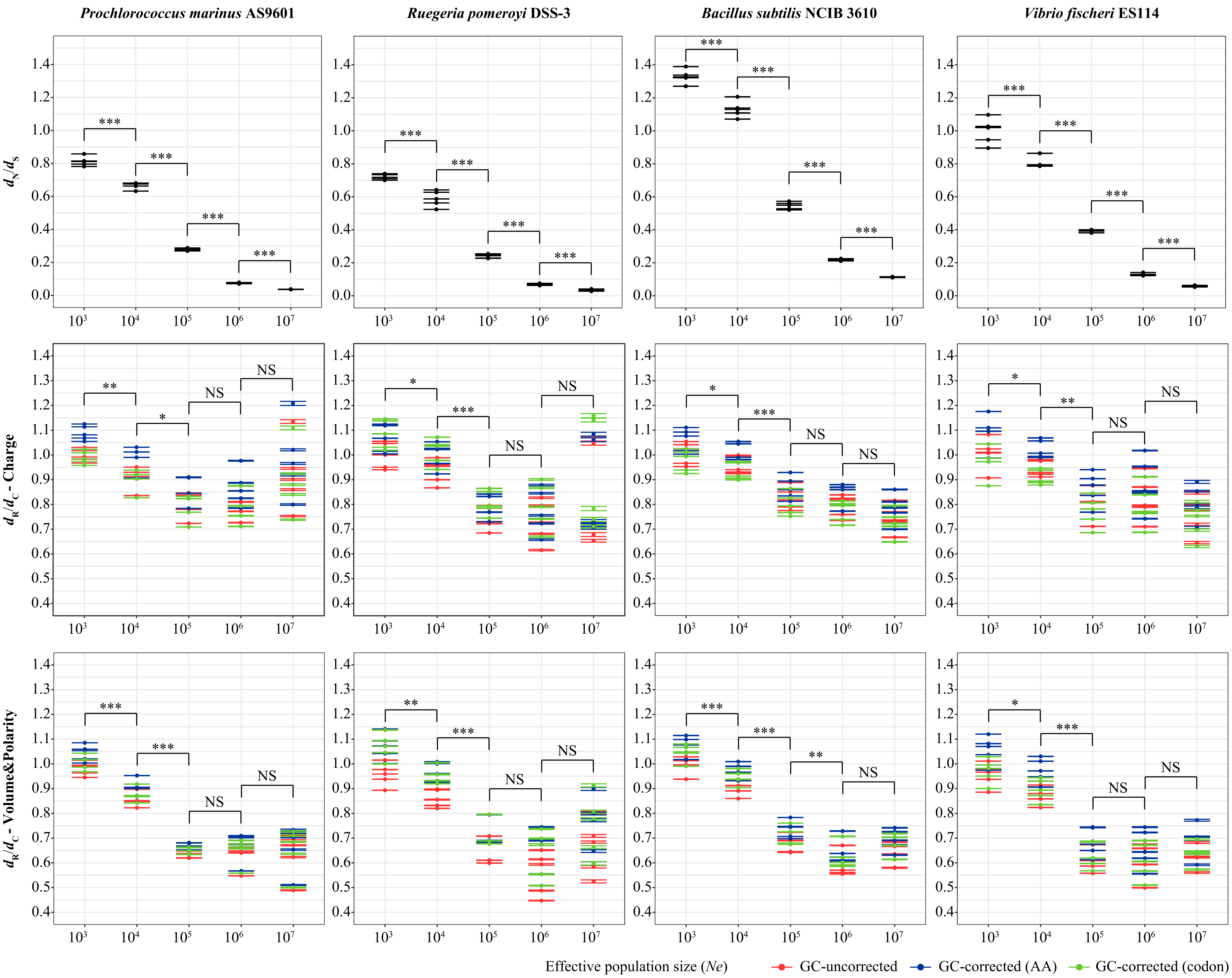


Fig. S3 The genome-wide $d_N/d_S$ and $d_R/d_C$ ratios of simulated populations of *Bacillus subtilis* BSn5, *Ruegeria pomeroyi* DSS-3, and *Vibrio fischeri* ES114. For each $N_e$, the simulation ran five times and each generated 50 genome sequences. Mean values of the $d_N/d_S$ and $d_R/d_C$ in each run are used for one-tailed t-test (***: $p<0.001$, **: $p<0.01$, *: $p<0.05$; NS: non-significant). The GC-corrected $d_R/d_C$ values based on codon frequency and AA composition are marked in red and blue, and the uncorrected $d_R/d_C$ values are marked in green. The error bars in the plots represent the standard error of the mean (SEM).

Fig. S4

A

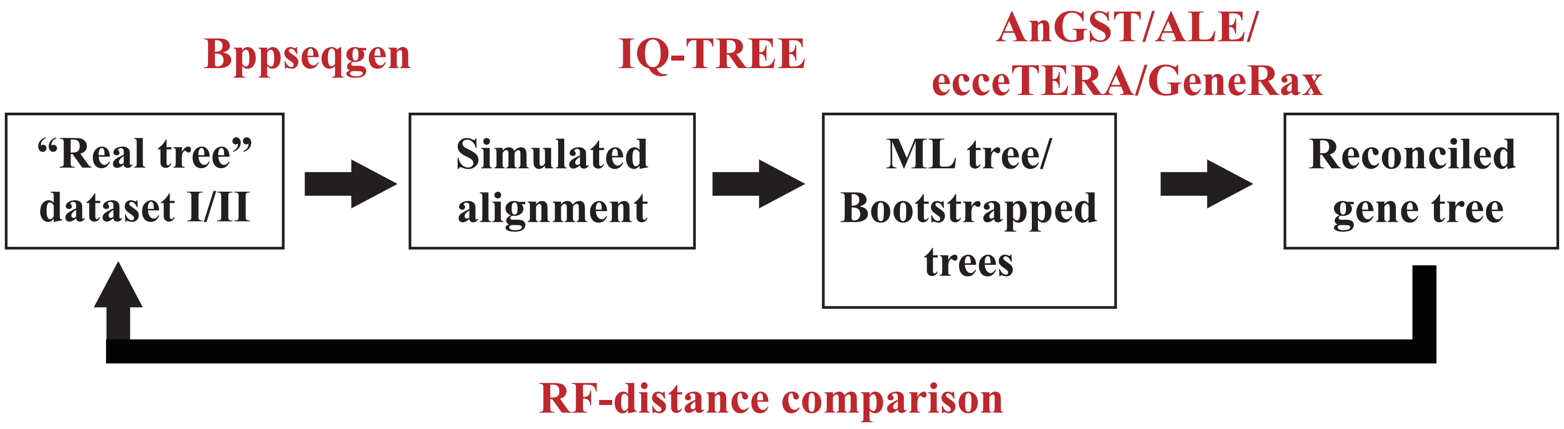


B

| RF distance | | "Real tree" dataset I | | "Real tree" dataset II | |
|---|---|---|---|---|---|
| | | Mean | SD | Mean | SD |
| ML | ALE | 0.2895 | 0.0888 | 0.3452 | 0.0806 |
| | GeneRax | 0.2659 | 0.0837 | 0.3319 | 0.1582 |
| MP | AnGST | 0.3457 | 0.1326 | 0.4204 | 0.0935 |
| | ecceTERA | 0.4024 | 0.1674 | 0.4761 | 0.1214 |

C

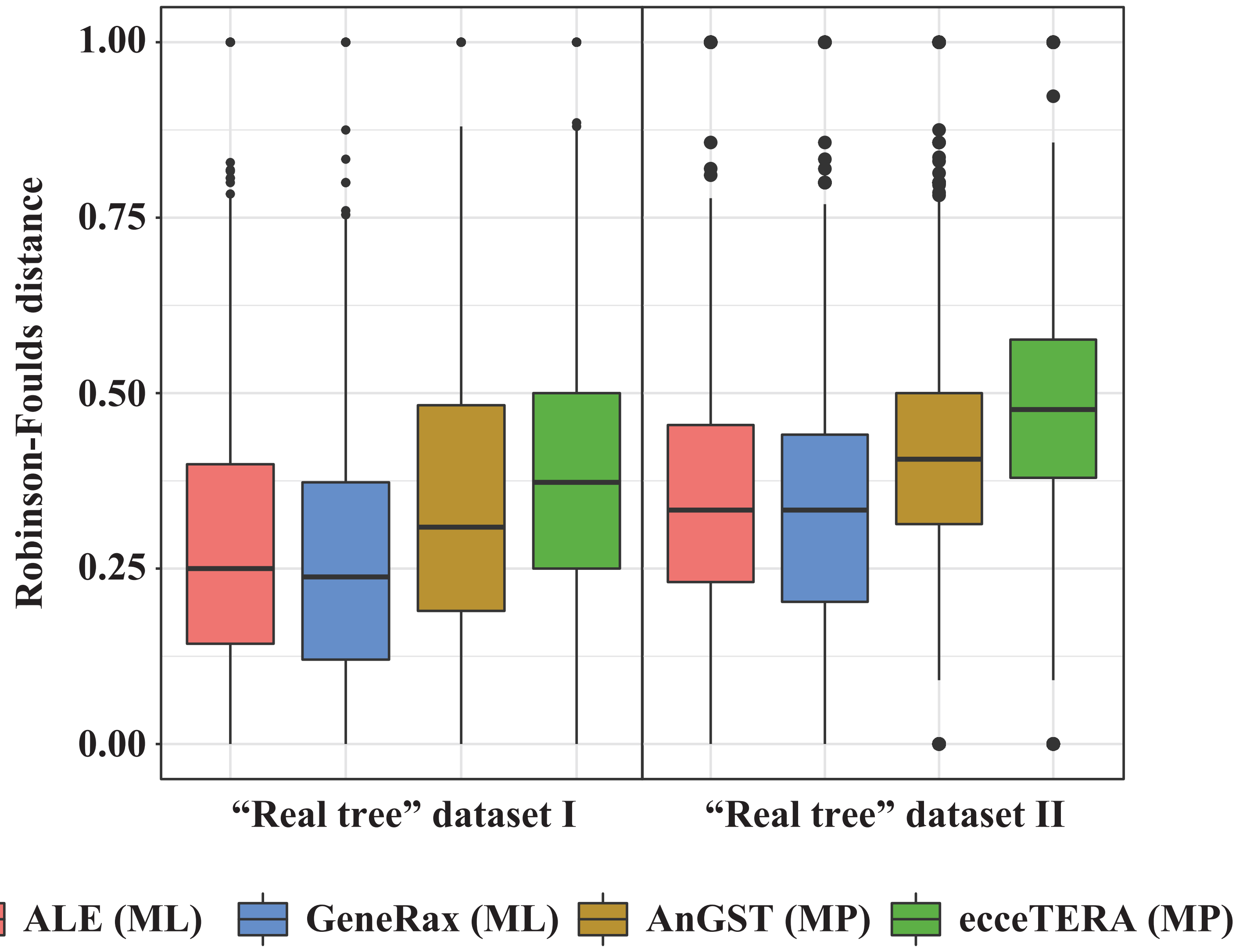


Fig. S4 (A) The illustration of the simulation-based benchmarking workflow, in which the tools are marked in red. (B) The mean and the standard deviation (SD) of the RF distance calculated based on the four reconciliation tools. MP: maximum parsimony-based reconciliation approach; ML: maximum likelihood-based reconciliation approach. The lower RF distance, the higher accuracy of the reconciliation. (C) The boxplot shows the RF distance calculated based on different tools and datasets. The middle line represents the median value of the RF distance. The lower and upper boundary of the box represent the 25th percentile and 75th percentile value of the RF distance, respectively. The upper and lower flank boundary represent the maximum and minimum RF distance, respectively, within the range of median ± 1.5 IQR (interquartile range: 75th percentile value - 25th percentile value). All other RF distance values, which are not included in this range, are marked with dots.
